# Y-box protein 1 is required to sort microRNAs into exosomes in cells and in a cell-free reaction

**DOI:** 10.1101/040238

**Authors:** Matthew Shurtleff, Kate V. Karfilis, Morayma Temoche-Diaz, Sayaka Ri, Randy Schekman

## Abstract

Exosomes are small vesicles that are secreted from metazoan cells and may convey selected membrane proteins and small RNAs to target cells for the control of cell migration, development and metastasis. To study the mechanisms of RNA packaging into exosomes, we devised a purification scheme based on the membrane marker CD63 to isolate a single exosome species secreted from HEK293T cells. Using immunoisolated CD63-containing exosomes we identified a set of microRNAs that are highly enriched with respect to their cellular levels. To explore the biochemical requirements for exosome biogenesis and RNA packaging, we devised a cell-free reaction that recapitulates the species-selective enclosure of miR-223 in isolated membranes supplemented with cytosol. We found that the RNA-binding protein Y-box protein I (YBX1) binds to and is required for the sorting of miR-223 in the cell-free reaction. Furthermore, YBX1 serves an important role in the secretion of miRNAs in exosomes by HEK293T cells.

## Introduction

In contrast to the normal pathways of protein secretion, the processes by which unconventional cargoes are secreted have proved diverse and enigmatic. Indeed, our understanding of unconventional secretory mechanisms is limited to a few examples of leader-less soluble and transmembrane proteins (Malhotra, 2013). Unconventionally secreted molecules may be externalized in a soluble form by translocation across various membranes. This may include direct translocation across the plasma membrane, or across an organelle membrane followed by fusion of the organelle with the plasma membrane (Zhang & Schekman, 2013). Alternatively, proteins and RNAs can be secreted within vesicles that bud from the plasma membrane, as in the budding of enveloped viruses such as HIV, or within vesicles internalized into a multivesicular body (MVB) that fuses with the plasma membrane (Colombo, Raposo, & Thery, 2014).

RNA is actively secreted into the medium of cultured cells and can be found in all bodily fluids enclosed within vesicles or bound up in ribonucleoprotin complexes, both forms of which are resistant to exogenous ribonuclease (Arroyo et al., 2011; Colombo et al., 2014; Mitchell et al., 2008). Importantly, extracellular vesicle-bound RNAs appear to be enriched in specific classes of RNAs, including small RNAs and microRNAs (Kosaka et al., 2010; Skog et al., 2008; Valadi et al., 2007).

Exosomes are a subclass of extracellular vesicle which can be defined as 30-100 |μm vesicles with a buoyant density of ∽1.10-1.19 g/ml that are enriched in specific biochemical markers, including tetraspanin proteins (Colombo et al., 2014). It is often assumed that vesicles fitting this description are derived from the multivesicular body, but some evidence suggests that physically and biochemically indistinguishable vesicles bud directly from the plasma membrane (Booth et al., 2006). Numerous studies have reported the presence of RNAs, especially microRNAs, from fractions containing exosomes, though many of these studies have relied on isolation techniques (e.g. high speed sedimentation) that do not resolve vesicles from other cellular debris or RNPs (Bobrie, Colombo, Krumeich, Raposo, & Thery, 2012). Thus, it is difficult to know in which form RNAs are secreted and even more challenging to determine which miRNAs may be specifically secreted as exosome cargo. The use of many different cell lines, bodily fluids and isolation methods to identify which miRNAs are specifically packaged into exosomes further complicates the establishment of widely accepted exosomal miRNA cargo.

Even with the crude preparations that have been characterized, it is clear that RNA profiles from exosomes are distinct from those of the producer cells. Thus RNA capture or stabilization in exosomes is likely to occur through a selective sorting mechanism. RNA packaging may occur by specific interactions with RNA binding proteins that engage the machinery necessary for membrane invagination into the interior of an MVB or by interaction of RNAs with lipid raft microdomains from which exosomes may be derived (Janas, Janas, Sapon, & Janas, 2015).

In order to probe the mechanism of exosome biogenesis, we developed procedures to refine the analysis of RNA sorting into exosomes. Using traditional means of membrane fractionation and immunoisolation, we identified unique miRNAs highly enriched in exosomes marked by their content of CD63. This miRNA sorting process was then reproduced with a cell-free reaction reconstituted to measure the packaging of exosome-specific miRNAs into vesicles formed in incubations containing crude membrane and cytosol fractions. Among the requirements for miRNA sorting in vitro, we found one RNA-binding protein, YBX1, which is a known constituent of exosomes secreted from intact cells (Buschow et al., 2010; Ung, Madsen, Hellwinkel, Lencioni, & Graner, 2014).

## Results

### Purified exosomes contain RNA

We first sought to purify exosomes from other extracellular vesicles and contaminating particles containing RNA (e.g. aggregates, ribonucleoprotein complexes) that sediment at high speed. We define exosomes as ∽30-100 nm vesicles with a density of 1.08-1.18 g/ml and containing the tetraspanin protein CD63. Based on these criteria, purified exosomes were recovered using a three-stage purification procedure (Figure 1a). First, large contaminating cellular debris was removed during low and medium speed centrifugation and exosomes were concentrated by high-speed sedimentation from conditioned medium. Next, to eliminate non-vesicle contaminants, the high-speed pellet fraction was suspended in 60% sucrose buffer and overlaid with layers of lower concentrations of sucrose buffer followed by centrifugation to float vesicles to an interface between 20 and 40% sucrose. Analysis of this partially purified material by electron microscopy showed vesicles of the expected size and morphology with fewer profiles of larger (>200 nm) membranes and reduced appearance of protein aggregates (Figure 1b,c compared to 1d,e). Finally, sucrose gradient fractions were mixed with CD63 antibody-immobilized beads to recover vesicles enriched in this exosome marker protein.

**Figure 1:**
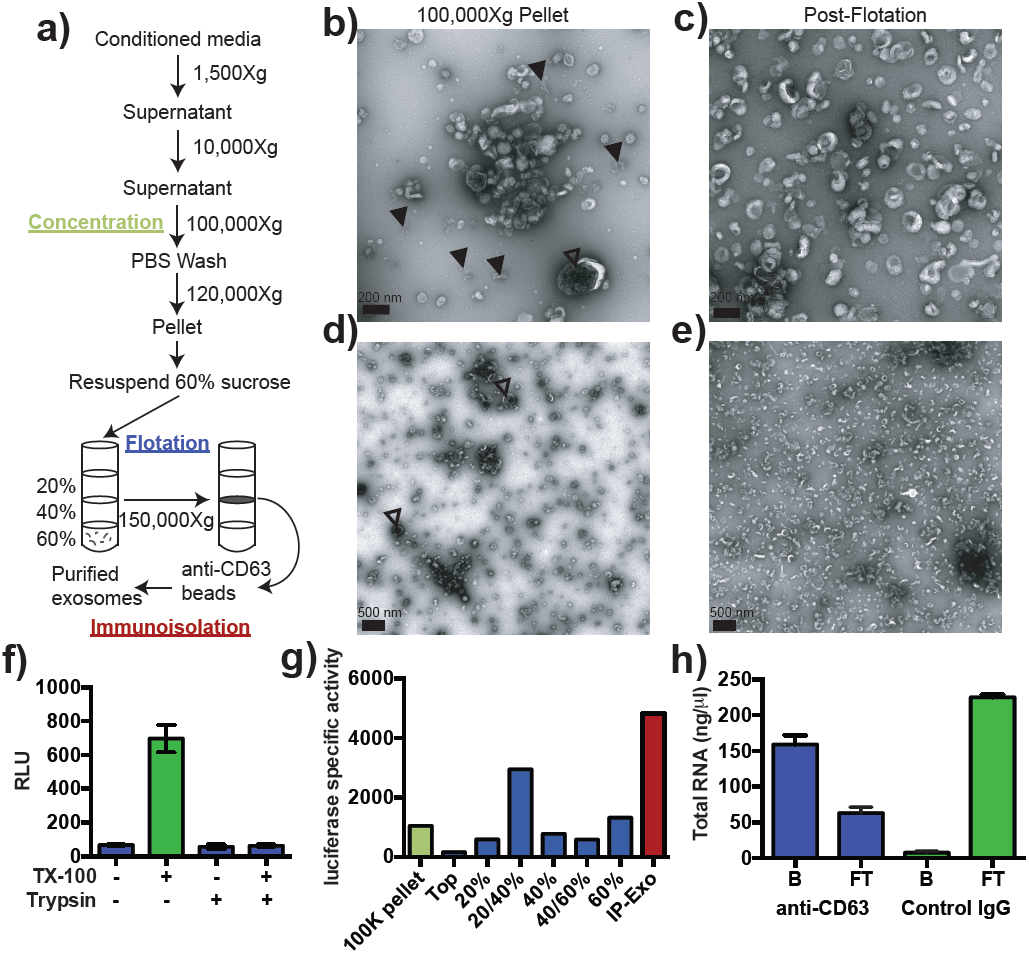
Purified CD63-positive exosomes contain RNA. **a)** Exosome purification schematic. **b-d)** Representative electron micrographs of negative stained samples from the 100,000Xg pellet fraction (b,d) and post-flotation fractions (c,e) at either 9,300X (a,b) or 1,900X (c,e) magnification. Open arrows indicate large (>200 nm) vesicle contaminants and closed arrows indicate protein aggregates. **f)** CD63‐ luciferase activity in purified exosomes after treatment with 1% Triton X-100 (TX-100) and/or 100 μg/ml trypsin for 30 min at 4°C. Error bars represent standard deviations from 3 independent samples. **g)** Specific activity of CD63-luciferase (RLU/μg of total protein) at each stage of purification (green: 100,000Xg pellet, purple: post-flotation, red: post-immunoisolation α-CD63 beads). **h)** Total RNA recovered from conditioned medium after immuno-isolation with α-CD63 or an IgG control. B – bound to beads, FT – flowthrough not bound to beads. Error bars represent standard deviations from 3 separate purifications.

To monitor and quantify the exosome purification, we generated a stable, inducible HEK293 cell line expressing a CD63-luciferase fusion. Tetraspanin proteins share a common topology in which the amino‐ and carboxyl-termini face the cytoplasm resulting in a predicted orientation inside the lumen of an exosomal vesicle. Although an intact C-terminal sequence is reported to be required for the proper localization of CD63 to the cell surface (Rous et al., 2002), we found that our overexpressed CD63-luciferase fusion was localized to a variety of cell surface and intracellular membranes (Figure 1‐ figure supplement 1). Using isolated exosome fractions, we confirmed that the CD63‐ luciferase fusion maintained the expected topology. Luciferase activity was stimulated by the addition of detergent to disrupt the membrane and allow access to the membrane impermeable substrates luciferin and ATP, and to trypsin, which inactivated luciferase activity in the presence but not in the absence of detergent (Figure 1f). The CD63‐ luciferase cell line was then used to monitor exosome purification. CD63-luciferase specific activity increased at each step of the purification, yielding a 5-fold purification of exosomes from the starting 100,000Xg pellet (Figure 1e). Additionally, following immunoisolation, most of the RNA was found in the CD63 positive bound (B) fraction, showing that RNA is associated with purified exosomes from HEK293T cells (1f). These results established that the RNA is associated with CD63-containing exosomes, but not necessarily enclosed within exosomes.

### Exosomes contain selectively packaged microRNAs

Previous reports indicated the presence of microRNAs (miRNAs) in fractions containing exosomes but which also contain contaminating particles (Bobrie et al., 2012; Skog et al., 2008; Valadi et al., 2007). To identify the specific miRNAs that are enriched in CD63‐ positive exosomes from 293T cells, we performed Illumina-based small RNA sequencing on libraries prepared from purified exosomes and from cells. We obtained a total of 58,848 miRNA reads in the exosome library representing 444 distinct miRNAs and 511,555 reads representing 549 miRNAs in the cell library (Figure 2a). To determine if a particular miRNA species was over-represented in exosomes, we analyzed the datasets for reads mapping to miRNA precursors and the targeting or passenger strand of mature miRNAs (Figure 2b). Exosomes were slightly enriched in reads mapping to precursor and passenger strand transcripts, however, the vast majority of miRNAs (91% from cells and 88% from exosomes) mapped to the mature targeting strand. The relative abundance of each miRNA was estimated by normalizing to the total number of miRNA-mapped reads (i.e. the number of reads mapping to a miRNA locus divided by the total number of mapped reads for each dataset). Of these, 56 and 161 miRNAs were uniquely found in the exosome and cell datasets respectively. Most of the miRNAs uniquely found in exosomes were of very low abundance, with only a few counts for each miRNA. The notable exception was miR-223, which was in the 72nd percentile for RPM in exosomes. The relatively high abundance of miR-223 in exosomes and its absence from the cellular library indicated that miR-223 was very efficiently packaged and secreted via exosomes (Figure 2c).

**Figure 2.**
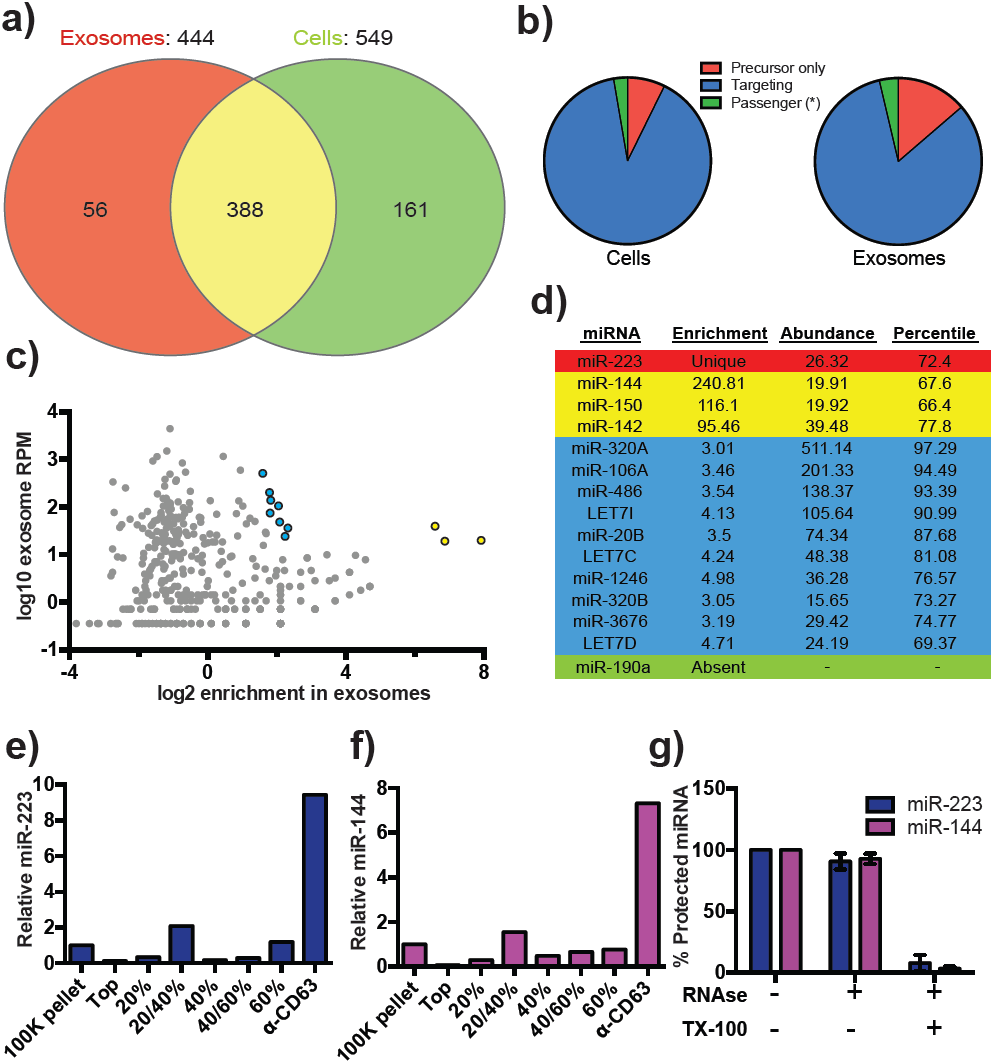
Enrichment of select microRNAs in exosomes. **a)** Venn diagram showing the number of total (above diagram), unique (inside red or green circles) and shared miRNAs (inside yellow) from each library. **c)** Scatterplot showing the enrichment (x-axis) and average relative abundance (y-axis) of all miRNAs found in both libraries. (Yellow markers – highly enriched miRNAs, Blue markers – moderately enriched miRNAs.) **d)** Table showing the enrichment, abundance (RPM) and percentile rank of relevant miRNAs in exosomes. (Red – unique to exosomes, Yellow – highly enriched in exosomes, Blue – moderately enriched, Green – unique to cells) **(e,f)** Pie charts showing the relative proportion of reads mapping to each miRNA species (Precursor only – red, passenger strand – green, targeting strand – blue) in cellular (e) and exosome (f) small RNA libraries **(g,h)** Relative miR-223 (e) and miR-144 (f) per ng of RNA as quantified by qRT-PCR during each stage of the purification. The 100K pellet was set to 1. **i)** RNase protection of exosomal miRNAs quantifed by qRT-PCR. Purified exosomes treated with or without RNase I and/or Triton X-100. Errors bars represent the standard deviation from 3 independent experiments.

We also identified miRNAs that were found in both libraries but were highly enriched in exosomes. A total of 388 miRNAs were detected in both the exosome and cell libraries. Most of these miRNAs were more enriched in cells than exosomes; however, miR144, miR150 and miR142 were highly enriched in exosomes and, like miR223, were moderately abundant in exosomes (Figure 2c and Figure 2d – yellow). In addition to very highly enriched miRNAs in exosomes, we identified moderately enriched (3-5 fold) miRNAs ranging from moderate to very high abundance in exosomes (69-97 percentile – blue). This cluster contained previously identified miRNAs enriched in exosomes, including highly abundant miR-320a and members of the Let7 family (Figure 2c, 2d – blue)(Guduric-Fuchs et al., 2012; Squadrito et al., 2014). To summarize, small RNA sequencing from purified exosomes and subsequent miRNA analysis identified several putative exosomal miRNAs.

### miR-223 and miR-144 are selectively packaged exosomal miRNAs

We next sought to confirm miR-223 and miR-144 as *bona fide* exosomal miRNAs. We performed quantitative reverse transcriptase PCR (qRT-PCR) for each miRNA target during the course of exosome purification from conditioned medium. Our results showed a selective enrichment of both miR-223 and miR-144 at each stage of the purification (Figures 2e and 2f). Thus, these miRNAs are associated with CD63 exosomes. To determine if these exosome associated RNAs are contained within exosomes and not simply bound to the surface, we performed an RNase protection experiment. Both miRNAs were protected from RNase I digestion, unless detergent was added to disrupt the membrane (Figure 2g). These results confirm that miRNAs are selectively packaged into exosomes purified from HEK293T conditioned media and establish miR-223 and miR-144 as specific exosomal miRNAs.

### Cell-free assays for exosome biogenesis and miRNA packaging

#### Rationale

The mechanism of exosome biogenesis has been probed in mammalian cell culture using the tools of gene knockdown, knockout and overexpression where it is often difficult to distinguish a primary or indirect role for a gene product. We sought to minimize these challenges by developing simple biochemical assays that reproduce an aspect of exosome biogenesis and miRNA packing in a cell-free reaction.

#### Exosome biogenesis in vitro

Since packaging of cargo into newly-formed vesicles presumably occurs concurrently with membrane budding, we sought to develop an assay to monitor the incorporation of an exosome membrane cargo protein into a detergent sensitive membrane formed in an incubation containing membranes and cytsolic proteins obtained from HEK293 cells (Figure 3a). Previous studies have reported the use of cell-free assays to monitor multivesicular body biogenesis and sorting of ubiquitinated membrane proteins into intraluminal vesicles (Falguieres et al., 2008; Tran, Chen, Emr, & Schekman, 2009; Wollert & Hurley, 2010). To specifically monitor exosome biogenesis, we measured the protection over time of luciferase fused to CD63. The fusion protein displayed luciferase on the cytoplasmic face of a membrane such that its incorporation into a vesicle by budding into the interior of an endosome (or into a vesicle that buds from the cell surface) would render the enzyme sequestered and inaccessible to exogenous luciferin and ATP, the substrates of catalysis (Fig. 1f). During the incubation, substrate would have access to luciferase exposed on the cytoplasmic face of a membrane or to enzyme about to be internalized into a bud, but not to luciferase that had already become sequestered within vesicles in cells prior to rupture (Fig. 3a). In order to focus only on luciferase that became sequestered during the cell-free reaction, aliquots taken after a 20 min incubation of membranes, cytosol and substrate at 30°C were sedimented, resuspended in buffer and sedimented again to remove excess substrate, and CD63-luciferase remaining exposed on the surface of membranes was inactivated by treatment with trypsin (0.5mg/ml) for 1h at 4°C. Remaining luciferase activity was then monitored in a luminometer. Since the substrates D-luciferin salt and ATP are membrane impermeable, the residual luminescence measured in trypsin-treated samples should derive from luciferase that became segregated along with substrate during the cell-free incubation. Relative protection was quantified as the ratio of relative light units (RLU) for reactions incubated without cytosol, at 4°C and in the presence of detergent (Triton X-100) divided by the RLU for a complete reaction incubated at 30°C. We observed that the formation of sequestered luciferase required cytosol and incubation at 30°C and was disrupted in incubations containing detergent (Triton X-100) (Figure 3b).

**Figure 3:**
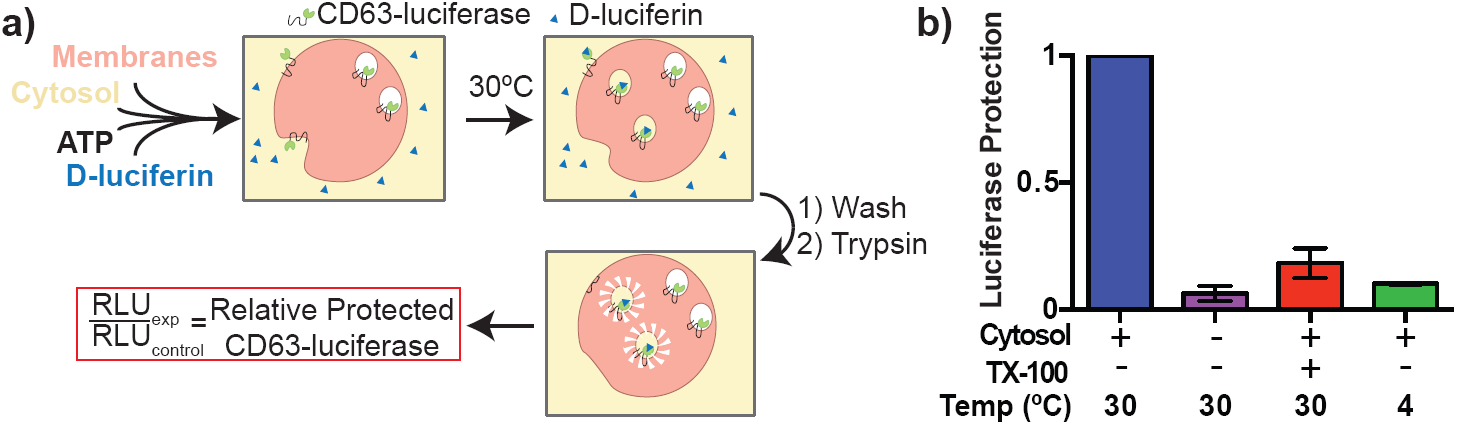
Cell-free exosome biogenesis reaction. **a)** Schematic illustrating the *in vitro* biogenesis reaction. **b)** Exosome biogenesis measured by relative protected CD63-luciferase. Reactions with or without cytosol, 1%Tx-100 and incubation temperature are indicated.

To examine the connection between the formation of sequestered luciferase and exosome biogenesis, we performed our cell-free reaction in the presence of an inhibitor (GW4869) of neutral sphingomyelinase, an enzyme that cleaves sphingolipid to form ceremide (Luberto et al., 2002). Treatment of cells with an inhibitor of neutral sphingomyelinase 2 (NS2) reduces the secretion of exosomes and exosome-associated miRNAs (J. Li et al., 2013; Trajkovic et al., 2008; Yuyama, Sun, Mitsutake, & Igarashi, 2012). GW4869 inhibited the protection of CD63-luciferase protection in our cell-free assay at concentrations required to inhibit NS2 activity in partially purified fractions of the enzyme (Figure 4d) (Luberto et al., 2002). Thus, our cell-free reaction may recapitulate an aspect of exosome biogenesis.

**Figure 4:**
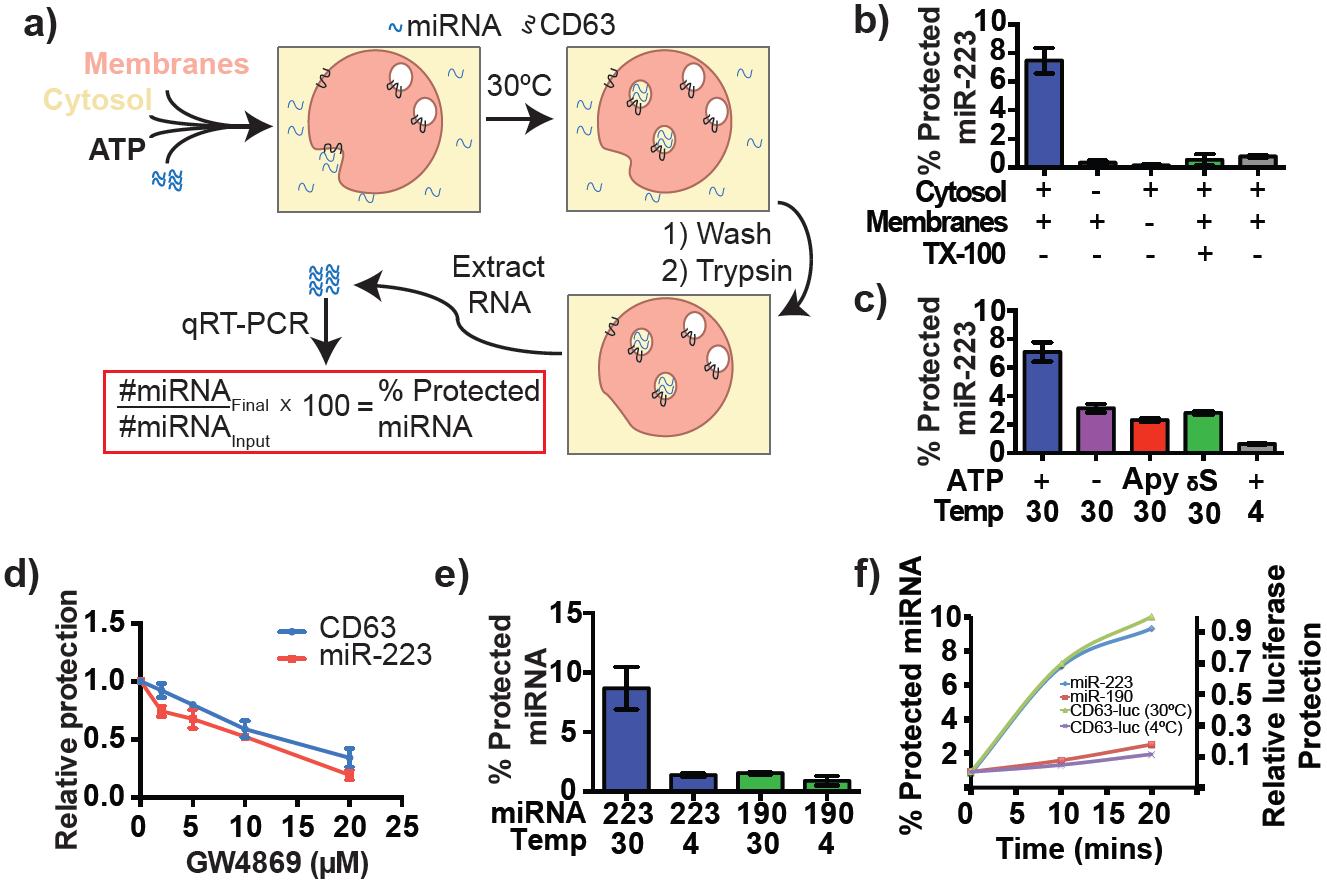
Cell-free selective sorting of miRNA into exosomes. **a)** Schematic illustrating the *in vitro* packaging reaction. **b)** Cell-free packaging of miR‐ 223 measured as percent protected by qRT-PCR. Reactions with or without membranes (15,000Xg pellet), cytosol (100,000Xg supernatant) and 1% Triton X-100 (TX-100), and incubated at 4 or 30 C are indicated. **c)** ATP requirements for miR-223 packaging (Apy – Apyrase, (1 U/ml) yS – ATPyS (10 mM)). **d)** Dose dependent effect of neutral sphingomyelinase 2 inhibitor (GW4869). Measured as relative protection of miRNA and CD63-luciferase normalized to vehicle only control (DMSO). **e)** Protection of miR-223 or miR-190 measured as a percent protected by qRT-PCR. Errors bars represent standard deviations from duplicate reactions. **f)** Relative CD63-luciferase (right axis) and percent miRNA protection (left axis) measured over a 20-min time course using the indicated miRNA cargo and incubation temperature.

#### MicroRNA packaging into vesicles in the cell-free reaction

Having identified miRNAs that are selectively packaged into exosomes in cultured cells, we examined our cell-free reaction for the RNA-selective segregation of miRNAs into RNase resistant and detergent sensitive vesicles (Figure 4a). As in the biogenesis reaction described above, crude membranes and cytosol from broken cells were mixed in buffer containing an ATP regenerating system, but in this case supplemented with synthetic miRNA, specifically miR-223, the miRNA that was most highly enriched in exosomes isolated from the medium of cultured HEK293T cells (Fig. 2d). Given the low relative abundance of miR-223 in HEK293T cells, we set the exogenous concentration in the incubation to be ca. 1000 fold in excess to ensure that the chemically synthetic material predominated in any packaged signal. After a 20 min incubation at 30°C, aliquots were treated with RNase I to digest any unpackaged miRNA. RNA was then purified, reverse transcribed using a miRNA specific primer and the amount of miRNA that became protected during the incubation was measured by quantitative PCR.

Packaged RNA was quantified as the percentage of miR-223 RNA molecules protected from RNase during the course of incubation. Packaging of miR-223 required membranes, cytosol and incubation at physiologic temperature (Figure 4b). As expected for the segregation of miR-223 into a membrane bound compartment, addition of TX-100 during the RNase incubation abrogated protection (Figure 4b). Furthermore, at a minimal concentration of cytosol (0.5 mg/ml), protection was stimulated two-fold over reactions performed in the in the absence of ATP, or in the presence of apyrase or a non-hydrolyzable analog of ATP (Figure 4c). The sphingomyelinase inhibitor GW4869 inhibited miR-223 packaging to the same extent and at similar concentrations to its effect on the formation of sequestered luciferase in the biogenesis reaction (Fig. 4d). Thus, the biogenesis and miRNA packaging reactions display similar biochemical requirements and may reflect the same process of exosome formation in our cell-free reaction.

### MicroRNA-223 is selectively packaged into vesicles *in vitro*

Having shown that specific miRNAs are enriched in exosomes produced *in vivo*, we next determined if selective sorting of miRNAs into vesicles could be reconstituted *in vitro.* We used the cell-free packaging assay to compare the efficiency of incorporation of synthetic miR-223 and a relatively abundant cellular miRNA that is not found in exosomes (miR-190). Exosomal miR-223 was more efficiently packaged into vesicles (9%) than cellular miR-190 (1.5%) (Figure 4e). Furthermore, the rate of miR-223 packaging mirrored the rate at which luciferase became sequestered in the biogenesis reaction whereas the rate of miR-190 protection in a 30°C incubation reflected the low rate of formation of sequestered luciferase in an incubation held on ice (Figure 4f). Based on these experiments, we conclude that the cell-free packaging assay reconstitutes the selective sorting of exosomal miR-223 over cellular miR-190 into vesicles, possibly exosomes, formed *in vitro*.

### Identifying candidate proteins involved in miRNA sorting into exosomes

To identify proteins that may be involved in miRNA packaging into exosomes, we employed a proteomics approach utilizing the *in vitro* packaging assay to capture RNA binding proteins. MiRNA sorting may require an RNA binding protein to segregate an RNP into a nascent budded vesicle. Synthetic 3’ biotinylated miR-223 was substituted for unmodified miRNA in the cell-free reaction. Samples were treated with RNase, quenched with RNase inhibitor and solubilized with Triton X-100. miR-223-biotin was captured on streptavidin-coated beads and interacting proteins were eluted with high salt buffer. Mir-223-interacting proteins were identified by in-solution liquid chromatography/mass spectrometry (Fig. 5a). Based on peptide count and coverage, the most highly represented protein was Y-box binding protein I (YBX1) (Fig. 5b). Peptides representing >45% YBX1 of the protein were identified stretching from the cold shock domain to the C-terminus (Fig. 5c).

YBX1 is a multi-functional RNA binding protein that shuttles between the nucleus, where it plays a role and splice site selection (Wang et al., 2013; Wei et al., 2012), and the cytoplasm where it is required for the recruitment of RNAs into cytoplasmic ribonucleoprotein granules containing untranslated mRNAs and plays a role in mRNA stability (Lyabin, Eliseeva, & Ovchinnikov, 2014). YBX1 also co-localizes with cytoplasmic P-bodies containing members of the RISC complex, including GW182 which can be found in exosomes (Gallois-Montbrun et al., 2007; Goodier, Zhang, Vetter, & Kazazian, 2007). Interestingly, YBX1 is secreted in a form that resists trypsin in the absence but not in the presence of a non-ionic detergent (Triton X-100), consistent with a location in vesicles, perhaps exosomes (Frye et al., 2009; Rauen et al., 2009). Furthermore, YBX1 has been detected by mass spectrometry in isolated exosomes (Buschow et al., 2010; Ung et al., 2014). Because YBX1 is a known RNA binding protein, is secreted by cells in vesicles and physically interacts with miR-223 during the *in vitro* packaging assay, it met our criteria for a potential exosomal miRNA sorting factor.

**Figure 5:**
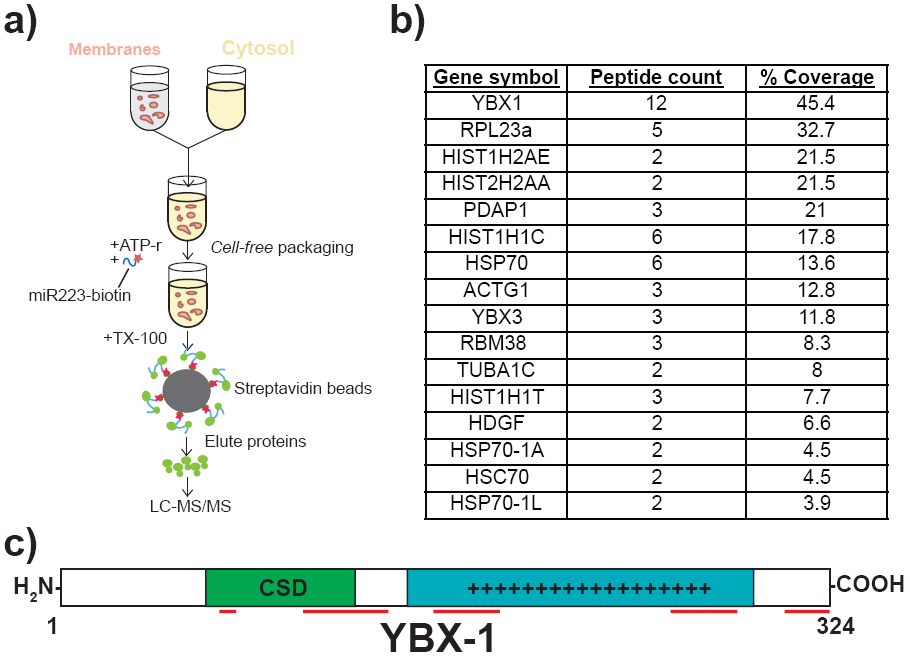
Identification of YBX1 as a candidate exosomal miRNA sorting protein. **a)** Scheme to identify candidate miRNA sorting proteins **b)** Proteins identified by tandem mass spectroscopy from the experiment illustrated in (a). **c)** Schematic of YBX1 protein. The cold-shock domain (green) and positively charged low-complexity region (blue) are highlighted. Red lines indicate detected unique peptides from mass spectroscopy.

### Y-box protein 1 is involved in sorting miR-223 into exosomes

We first determined if YBX1 co-purifies with exosomes. We purified exosomes as in Fig. 1a and found that YBX1 was primarily associated with the CD63-bound fraction containing known exosome markers (TSG101, Alix, CD9), as opposed to Flotillin 2, which was predominantly found in the unbound fraction (Fig. 6a). The CD63 positive (exosome) fraction contained most of the RNA (Fig. 1f). These results show that HEK293T cells release at least two vesicle types (CD63 positive and negative).

**Figure 6:**
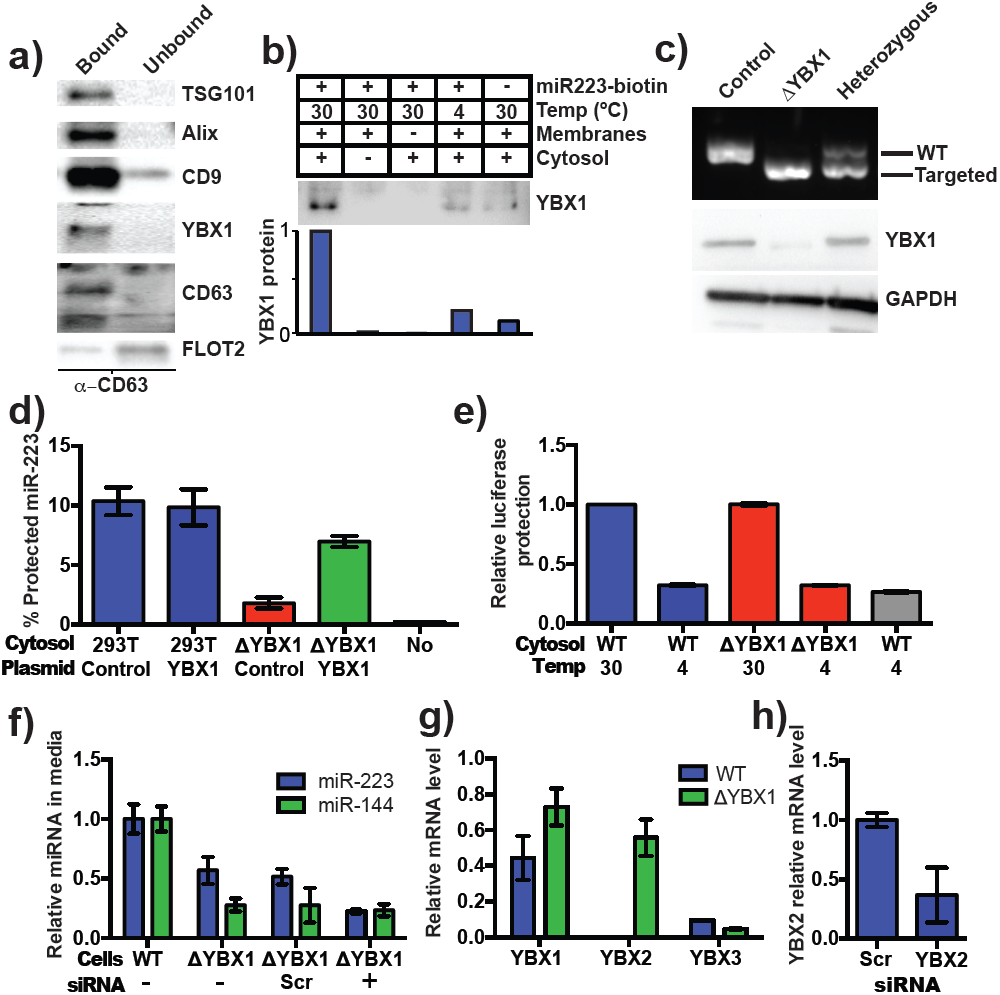
YBX1 is necessary for exosomal miRNA packaging and secretion. **a) Immuno**blots for the indicated protein markers in the CD63 immuno-isolated (bound) or unbound fractions. Exosomes were purified as in Figure 1a. **b) Immuno**blot for YBX1 following cell-free packaging reactions performed according to the conditions indicated and immobilized with streptavidin beads as shown in (Figure 5a). Bar graph represents densitometry values for the blot shown. **c)** Analysis of wild-type and CRISPR/Cas9 genome edited HEK293T clones by PCR flanking the genomic target site (top) and immunoblot for YBX1 (middle) and GAPDH (bottom). **d)** *In vitro* miR-223 packaging into exosomes from ΔYBX1 or WT cytosol transfected with control (pCAG) or YBX1 plasmid, **e)** Cell-free exosome biogenesis with cytosol from ΔYBX1 or WT cells and membranes from CD63-luciferase cells. **f)** Relative quantity of miR-223 secreted into the medium by WT and ΔYBX1 cells after 24 h with or without transfection with control or YBX2 siRNA. **h)** Relative mRNA expression of the human Y-box paralogs in WT and ΔYBX1 cells. Measured by qRT-PCR using the ΔΔCt method and normalized to GAPDH. **g)** YBX2 expression in ΔYBX1 cells after knockdown with YBX2 or negative control siRNA. Error bars represent standard deviations from triplicates in all panels.

We next examined the biochemical requirements for co-packaging of miR-223 and YBX1 using the biotin-miR-223 pull down assay described in Fig. 5a. An immunoblot showed YBX1 bound to biotin-mi223 was recovered in exosomes in a complete reaction, while no detectable YBX1 was recovered in exosomes in incubations that lacked cytosol or membranes and much reduced signals in control incubations held at 4C or conducted in the absence of biotin-mi223 (Fig. 6b). These conditions mirror those required for the packaging of miR-223 in our cell-free reaction.

We next evaluated the requirement for YBX1 in packaging exosomal miRNAs in cells and in the cell-free reaction. To address this question, we generated a YBX1 knockout HEK293T cell line with CRISPR/Cas9 using a guide RNA targeting the YBX1 locus (Cong et al., 2013; Jinek et al., 2013; Mali et al., 2013). Clones were screened by genomic PCR and immunoblot for YBX1. We recovered a homozygous mutant clone (ΔYBX1) that had been targeted at the YBX1 locus and no longer expressed YBX1 protein (Fig. 6c). The homozygous mutant cells grew normally under the conditions used to propagate HEK293T cells and released an approximately equal number of particles into the medium after 48 h of growth (2.38X10^7^ and 2.42X10^7^ particles/ml for wild type and ΔBX1) as determined by Nanosight nanoparticle tracking analysis.

To determine if YBX1 was required for miRNA packaging, we prepared cytosol from ΔYBX1 cells and tested miR-223 incorporation using the *in vitro* packaging assay. Cytosol from ΔYBX1 cells did not support miR-223 protection *in vitro* but activity was largely restored in reactions containing cytosol from a ΔYBX1 line transfected with plasmid encoding YBX1 (Fig. 6d). We also evaluated the role of YBX1 in our biogenesis reaction (Fig. 3a) and found that cytosol from wt and ΔYBX1 were indistinguishable in the formation of latent luciferase activity (Fig. 6e). Thus, YBX1 is required for exosomal miRNA packaging *in vitro* but is not required for the sorting of an exosome membrane cargo protein into vesicles in our cell-free reaction.

In an effort to connect the results of our cell-free reaction to the mechanism of sorting of miRNAs into exosomes secreted by HEK293T cells, we examined the secretion of miR-223 and of another miRNA, miR-144, which was also highly enriched in our purified exosome fraction (Fig. 2d). We measured the amount of miR-144 and – 223 secreted into the medium by qRT-PCR. ΔYBX1 cells showed a substantial decrease in secretion of miR-144 but only ∽50% decrease in miR-223 secretion during a 24 h incubation in fresh medium (Fig. 6f).

The residual secretion of miR-223 in ΔYBX1 cells may be due to partial redundancy conferred by other YBX1 paralogs (YBX2 and YBX3). YBX3 is constitutively expressed at a low level in most cell types whereas YBX2 expression is restricted to germ line cells (Gu et al., 1998). We measured the relative expression of mRNAs encoding all three Y-box paralogs in WT and ΔYBX1 cells and found that YBX2 mRNA expression was increased >500 fold in ΔYBX1 cells compared to WT (Fig. 6g). Thus, ΔYBX1 cells exhibited compensatory expression of the germ-line specific transcript, YBX2. When YBX2 expression was ablated by siRNA in ΔYBX1 cells, miR‐ 223 secretion was further diminished to the low level seen for miR-144 in ΔYBX1 cells (Fig. 6g,h). These data show that YBX1 is involved in exosomal miRNA (miR-144 and miR-223) secretion *in vivo.* The Y-box paralogs may play a similar role in miRNA secretion in other cell types.

## Discussion

Our results establish miR-223 and miR-144 as specific exosome cargo in HEK293T cells. In order to probe the mechanism of RNA sorting into exosomes, we developed biochemical assays that measure the capture of an exosome membrane protein and miRNA into vesicles formed in a cell-free reaction. Using this approach, we identify YBX1 as an RNA-binding protein that is critical for the efficient packaging of miR-223 *in vitro* and its secretion in cultured human cells.

How are miRNAs recognized for sorting into exosomes? We find that synthetic, miR-223 is sequestered into vesicles more efficiently than miR-190, consistent with the possibility of a primary RNA sequence or secondary structure, perhaps stabilized by an RNA binding protein such as YBX1, that directs RNA sorting. One possible sorting motif - GGAG - is enriched in miRNAs secreted in exosomes from T-cells (Villarroya‐Beltri et al., 2013). This motif is recognized by hRNPA2B1, a T-cell exosome RNA‐binding protein, which requires sumoylation for efficient secretion via exosomes. It was therefore suggested that binding of GGAG containing miRNAs by sumoylated hRNPA2B1 was a sorting mechanism for miRNAs into T-cell-derived exosomes.

We were unable to identify any statistically significant primary sequence motifs for miRNAs by either multiple alignment (ClustalW) or multiple Em for motif elicitation (MEME) in HEK293T-derived exosomes (Bailey et al., 2009; Larkin et al., 2007). Furthermore, the mature targeting strand of miR-223 packaged into exosomes contains no guanine nucleotides and hRNPA2B1 was not detected in our mass spectrometry results for proteins bound to miR-223-biotin isolated from vesicles formed in our cell-free reaction. The human genome encodes more than 1,000 experimentally determined and predicted RNA binding proteins (Cook, Kazan, Zuberi, Morris, & Hughes, 2011; Gerstberger, Hafner, & Tuschl, 2014). We therefore propose that different cell types may use RNA binding proteins with distinct binding preferences to secrete miRNAs, and perhaps other RNA classes, in exosomes. In addition, some cell types may deploy multiple RNA-binding proteins to sort RNAs into exosomes, in which case motif discovery would be challenging, even in highly purified vesicles, due to diverse motif preferences from distinct proteins.

We identified YBX1 as the dominant RNA-binding protein physically interacting with miR-223 *in vitro* and confirmed its role in miR-223 packaging into exosomes both *in vitro* and in cultured cells. YBX1 is found within mammalian P-bodies (GW bodies) containing untranslated RNAs (Kedersha & Anderson, 2007) and in the nucleus where it plays a role in RNA splice site selection by binding short sequence motifs (Wang et al., 2013; Wei et al., 2012). YBX1 binds RNA via an internal cold shock domain and an inherently disordered, highly charged C-terminus (Lyabin et al., 2014). Interestingly, another cold shock domain containing protein, Lin28, binds pre-miRNAs of the Let7 family via hairpin-loop structures (Nam, Chen, Gregory, Chou, & Sliz, 2011). YBX1 also binds hairpin-loops in a murine retrovirus, leading to stabilization of the viral RNA genome and increased particle production (Bann, Beyer, & Parent, 2014). YBX1 binding of viral RNA also increases production of other retroviruses, including HIV (Bann et al., 2014; W. Li, Wang, & Gao, 2012; Mu, Li, Wang, & Gao, 2013). This raises the possibility that the recognition motif for sorting into exosomes may be based on secondary rather than primary RNA structure and that YBX1 may act as an RNA cofactor to escort exosomal RNAs into exosomes.

Our studies focused on miRNAs, however it is possible that YBX1 is responsible for the secretion of other RNA classes in exosomes. Several recent reports indicate a role for YBX1 in binding various small RNAs, including miRNAs, tRNA fragments and snoRNAs (Blenkiron, Hurley, Fitzgerald, Print, & Lasham, 2013; Goodarzi et al., 2015; Liu et al., 2015). It is interesting to note that most miRNAs present in exosomes in our study are not highly enriched compared to their relative abundance in cells. This raises the possibility that highly enriched exosomal miRNAs mimic other classes of RNAs that are more efficiently packaged in a YBX1-dependent manner.

YBX1 secretion requires two acetylated lysine residues in the C-terminal domain (K304 and K307), as substitution of alanines for these lysines blocks YBX1 secretion (Frye et al., 2009). Interestingly, these residues are also ubiquitylated *in vivo* (Kim et al., 2011). Ubiquitylation of membrane proteins on the surface of an endosome targets them for internalization by the ESCRT machinery involved in the sorting and invagination of vesicles to form the multivesicular body. By analogy, ubiquitylation could direct YBX1 and its RNA cargo to the site of membrane invagination to form the precursor of an exosome. Indeed, several studies have reported the presence of ubiquitylated proteins in exosomes (Burke, Oei, Edwards, Ostrand-Rosenberg, & Fenselau, 2014; Buschow, Liefhebber, Wubbolts, & Stoorvogel, 2005). The exact role of acetylation, ubiquitylation and the ESCRT complexes in the sorting and secretion of proteins and RNA in exosomes remains an open question.

The functional role of secreted miRNAs has been a matter of discussion since the first reports of extracellular RNA (Thery, 2011). Numerous studies have shown that miRNAs can be transferred to neighboring cells in experimental settings (Cha et al., 2015; Kosaka et al., 2010; Mittelbrunn et al., 2011; Pegtel et al., 2010; Rana, Malinowska, & Zoller, 2013; Skog et al., 2008; Valadi et al., 2007). However, the transfer of miRNAs in biologically significant quantities for function in a physiological context is far from proven. Indeed, a recent study reported a stoichiometry of less than one specific miRNA per exosome, with the caveat that this study characterized crude, high-speed pellet fractions from conditioned medium (Chevillet et al., 2014). Functional miR-223 transferred between macrophages and miR-223-containing exosomes can induce macrophage differentiation, however, it has yet to be shown that miR-223 transfer plays a direct role in the differentiation (Ismail et al., 2013). Indeed, direct and convincing evidence for a physiological role of miRNAs secreted via exosomes has so far proven elusive.

Alternatively, exosomes may be a convenient carrier to purge unnecessary or inhibitory RNAs from cells. A recent report provided evidence for both alternative views with the demonstration that target transcript levels for miRNAs in the cell modulate the abundance of miRNAs in macrophage exosomes, and this in turn dictates which miRNAs are transferred to repress transcripts in recipient cells (Squadrito et al., 2014). Because YBX1 and the RISC machinery have both been shown to localize to P-bodies and P-bodies are closely juxtaposed to multivesicular bodies, all of the necessary machinery is poised to efficiently secrete miRNAs in exosomes (Gibbings, Ciaudo, Erhardt, & Voinnet, 2009). YBX1 may complex with miRNAs whose mRNA targets are not expressed, and sort them into the intralumenal vesicles of a multivesicular body for export by unconventional secretion.

## Materials and Methods

### Cell lines, media and general chemicals

HEK293T cells were cultured in DMEM with 10% FBS (Thermo Fisher Scientific, Waltham, MA). For exosome production, cells were seeded to ∽10% confluency in 150 mm CellBIND tissue culture dishes (Corning, Corning NY) containing 30 ml of growth medium and grown to 80% confluency (∽48 h). We noted that confluency >80% decreased the yield of exosome RNA. Cells grown for exosome production were incubated in exosome-free medium produced by ultracentrifugation at 100,000Xg (28,000 RPM) for 18h using an SW-28 rotor (Beckman Coulter, Brea, CA) in a LE-80 ultracentrifuge (Beckman Coulter). Unless otherwise noted, all chemicals were purchased from Sigma Aldrich (St. Louis, MO)

### Exosome purification

Conditioned medium (3 l for small RNA-seq and 420 ml for all other experiments) was harvested from 80% confluent HEK293T cultured cells.. All subsequent manipulations were performed at 4°C. Cells and large debris were removed by centrifugation in a Sorvall R6+ centrifuge (Thermo Fisher Scientific) at 1,500Xg for 20 min followed by 10,000Xg for 30 min in 500 ml vessels using a fixed angle FIBERlite F14-6X500y rotor (Thermo Fisher Scientific). The supernatant fraction was then passed through a 0.22 μM polystyrene vacuum filter (Corning) and centrifuged at ∽100,000Xg (26,500 RPM) for 1.5 h using two SW-28 rotors. The maximum rotor capacity was 210 ml, thus the small RNA-seq processing required pooling from ∽15 independent centrifugations. The pellet material was resuspended by adding 500 μl of phosphate buffered saline, pH 7.4 (PBS) to the pellet of each tube followed by trituration using a large bore pipette over a 30 min period at 4°C. The resuspended material was washed with ∽5 ml of PBS and centrifuged at ∽120,000Xg (36,500 RPM) in an SW-55 rotor (Beckman Coulter). Washed pellet material was then resuspended in 200 μl PBS as in the first centrifugation step and 1 ml of 60% sucrose buffer (20 mM Tris-HCl, pH 7.4 137 mM NaCl) was added and mixed with the use of a vortex to mix the sample evenly. The sucrose concentration in the PBS/sucrose mixture was measured by refractometry and, if necessary, additional 60% sucrose buffer as added until the concentration was >50%. Aliquots (1 ml) of 40%, 20% and 0% sucrose buffer were sequentially overlaid and the tubes were ultracentrifuged at ∽150,000Xg (38,500 RPM) for 16 h in an SW-55 rotor. The 20/40% interface was harvested, diluted 1:5 with phosphate buffered saline (pH 7.4) and 1 μg of rabbit polyclonal anti-CD63 H-193 (Santa Cruz Biotechnology, Dallas, TX) was added per liter of original conditioned medium and mixed by rotation for 2 h at 4C. Magvigen protein‐A/G conjugated magnetic beads (Nvigen, Sunnyvale, CA) were then added to the exosome/antibody mixture and mixed by rotation for 2 h at 4C. Beads with bound exosomes were washed three times in 1 ml PBS and RNA was extracted using Direct-Zol RNA mini-prep (Zymo Research, Irvine, CA) or protein was extracted in 100 μl 1X Laemmli sample buffer and dispersed with the use of a vortex mixer for 2 min.

### Negative staining and visualization of exosomes by electron microscopy

An aliquot (4 ul) of the resuspended 100,000Xg pellet fraction or a sample from the 20/40% interface that was diluted 10-fold with PBS, centrifuged at 100,000Xg in a TLS-55 rotor and then resuspended in 1 % glutaraldehyde, was spread onto glow discharged Formvar-coated copper mesh grids (Electron Microscopy Sciences, Hatfield, PA) and stained with 2% Uranyl acetate for 2 min. Excess staining solution was blotted off with filter paper. Post drying, grids were imaged at at 120 kV using a Tecnai 12 Transmission Electron Microscope (FEI, Hillsboro, OR) housed in the Electron Microscopy Laboratory at UC Berkeley.

### Nanoparticle Tracking Analysis

Conditioned medium (1 ml) from wild type and ΔYBX1 cells was harvested and the supernatant from a 10,000Xg centrifugation was drawn into a 1 ml syringe and inserted into a Nanosight LM10 instrument (Malvern, UK). Particles were tracked for 60 s using Nanosight nanoparticle tracking analysis software. Each sample was analyzed 4 times and the counts were averaged.

### Construction of inducible 293:CD63-luciferase cell line and luciferase activity assays

HEK293 cells expressing doxycycline-inducible CD63-luciferase was generated using the T-REx – 293 cell line according to the manufacturer’s instructions (Life Technologies, Grand Island, NY). The open reading frame for CD63-was amplified from human cell cDNA and firefly luciferase-FLAG was amplified from a plasmid source, both using Phusion DNA Polymerase (NEB). CD63 was fused to luciferase by NotI digestion, ligation and PCR amplification. The CD63-luciferase-FLAG amplicon was then digested and ligated into pcDNA5/FRT/TO (Life Technologies) using NdeI and PstI sites. The resulting plasmid was co-transfected with pOG44 (Life Technologies) and a stable cell line was selected using hygromycin selection (100 μg/ml). CD63-luciferase expression was induced with 1 μg/ml doxycycline 48 h prior to exosome harvesting. Luciferase activity was measured using a Promega Glowmax 20/20 luminometer (Promega, Madison, WI) with a signal collection integration time of 1 s. Luciferase reactions contained 50 μl sample, 10 μl 20X luciferase reaction buffer (500 mM Tricine, pH 7.8, 100 mM MgSO4, 2 mM EDTA), 10 μl 10 mM D-luciferin dissolved in PBS, 10 μl ATP dissolved in deionized water and 120 μl deionized water. Where indicated, samples were pre-treated with final concentrations of 1% Triton X-100 and/or 100 μg/ml trypsin for 30 min on ice. Total protein concentrations were measured using Pierce BCA protein assay according to the manufacturer’s instructions.

### Immunoblotting

Exosome and cell lysates were prepared by mixing in lysis buffer (10 mM Tris-HCl, pH 7.4, 100 mM NaCl, 0.1% sodium dodecyl sulfate, 0.5% sodium deoxycholate, 1% Triton X-100, 10% glycerol). Lysates were diluted 4-fold with 4X Laemmli sample buffer, heated to 65°C for 5 min and separated on 4-20% acrylamide Tris-Glycine gradient gels (Life Technologies). Proteins were transferred to polyvinylidene difluoride membranes (EMD Millipore, Darmstadt, Germany), blocked with 5% bovine serum albumin in TBST and incubated overnight with primary antibodies. Blots were then washed with TBST, incubated with anti-Rabbit or anti-Mouse secondary antibodies (GE Healthcare Life Sciences, Pittsbugh, PA) and detected with ECL-2 reagent (Thermo Fisher Scientific). Primary antibodies used in this study were anti-YBX1 (Cell Signaling Technology, Danvers, MA), anti-GAPDH (Santa Cruz Biotechnology), anti-TSG101 (Genetex, Irvine, CA), anti-CD9 (Santa Cruz Biotechnology), anti-Flotillin 2 (Abcam, Cambridge, MA), anti-Alix (Santa Cruz Biotechnology).

### Quantitative real-time PCR

RNA was extracted using the Direct-Zol RNA mini-prep and cDNA was synthesized either by oligo-dT priming (mRNA) or gene-specific priming (miRNA) according to the manufacturer’s instructions. For miRNA, we used Taqman miRNA assays from Life Technologies (assay numbers: hsa-mir-223-3p: 000526, hsa-mir-190a-5p: 000489 and hsa-miR-144-3p: 002676). Because there is no well-accepted endogenous control transcript for exosomes, relative quantification was performed from equal amounts of total RNA. Qubit (Thermo Fisher Scientific), was used to quantify total RNA from the medium or cells: 10 ng of RNA was reverse transcribed and qPCR was performed according to manufacturer’s instructions. Relative quantification was calculated from the expression 2^^^-(Ct_control_-Ct_experimental_). For mRNA quantification, total mRNA from cells was reverse transcribed using Superscript III enzyme and oligo-dT priming according to the manufacturer’s instructions (Life Technologies). qPCR was performed using SYBR green master mix (Life Technologies) using the primers: YBX1-F (CAACAGGAATGACACCAAGGA), YBX1-R (GTGTAGGAGATGGAGAGACTGT), YBX2-F (TCTTTGTTCACCAGACAGCTATTA), YBX2-R (CTCTCCTTCCACGACATCAAAT), YBX3-F (TGTACATCAGACTGCCATCAAG), YBX3-R (CCTTCTCTCCTTCAACCACATC), GAPDH-F (ACCCACTCCTCCACCTTTGAC), and GAPDH-R (TGTTGCTGTAGCCAAATTCGTT). Relative quantification was calculated using the 2^^-ΔΔ^Ct^^ method (Livak & Schmittgen, 2001). Taqman and SYBR green master mixes were obtained from Life Technologies and quantitative real-time PCR was performed using an ABI-7900 real-time PCR system (Life Technologies).

### Cell-free Biochemical Assays

#### Preparing membranes and cytosol

HEK293T cells were harvested at ∽80% confluency by gently pipetting with PBS. Cells were centrifuged at 500Xg and the pellet was weighed and frozen at −80C until use. Cells were thawed, resuspended in 2 vol of homogenization buffer (250 mM sorbitol, Tris-HCl, pH 7.4, 137 mM NaCl) containing protease inhibitor cocktail (1mM 4-aminobenzamidine dihydrochloride, 1 μg/ml antipain dihydrochloride, 1 μg/ml aprotinin, 1 μg/ml leupeptin, 1 μg/ml chymostatin, 1 mM phenymethylsulfonly fluoride, 50 μM N-tosyl-L-phenylalanine chloromethyl ketone and 1 μg/ml pepstatin) and passed 7-15 times through a 22 gauge needle until >80% of cells were disrupted, as assessed by microscopy and trypan blue staining. The homogenized cells were then centrifuged at 1,500Xg and the supernatant fraction was centrifuged at 15,000Xg using a FA-45-30-11 rotor and Eppendorf 5430 R centrifuge (Eppendorf, Hamburg, Germany). The supernatant fraction was centrifuged again at 55,000 RPM in a TLS-55 rotor and Optima Max XP ultracentrifuge (Beckman Coulter) to generate the cytosol fraction (∽5 mg/ml). The 15,000Xg pellet fraction was resuspended in 2 packed cell vol homogenization buffer and an equal vol of 1 M LiCl. The membranes were then centrifuged again at 15,000Xg and resuspended in 1 original packed cell vol to generate the membrane fraction.

#### Cell-free exosome biogenesis assay

Membranes were prepared from HEK293:CD63-luciferase cells and cytosol from HEK293T cells. Complete biogenesis reactions (40 μl) consisted of 10 μl membranes, 17 μl cytosol (2 mg/ml final concentration) + homogenization buffer, 4 μl 10X ATP regeneration system (10mM ATP, 500 mM GDP-mannose, 400 mM creatine phosphate, 2mg/ml creatine phosphokinase, 20 mM HEPES, pH 7.2, 250 mM sorbitol, 150 mM KOAc, 5 mM MgOAc), 8 ul 5X incorporation buffer (80 mM KCl, 20 mM CaCl2, 12.5 mM HEPES-NaOH, pH 7.4, 1.5 mM, MgOAc, 1 mM DTT), 1 μl D-luciferin (10 mM in PBS). The reaction mixture was incubated at 30°C for 20 min and membranes were sedimented at 15,000Xg at 4°C. Post-reaction membranes were then resuspended in 500 ul PBS with 0.5 mg/ml trypsin and incubated at 4°C for 1h to inactivate CD63-luciferase that had not been internalized during the incubation period. The trypsin-treated reactions were then incubated for 2 min at 25°C and luciferase activity was quantified using the luminometer conditions described above. Exosome biogenesis for experimental conditions was calculated as the relative ratio compared to the complete control reaction described above (RLU_experimental_/RLU_control_).

#### Cell-free exosome miRNA packaging assay

Membranes and cytosol were prepared from HEK293T cells. Complete miRNA packaging assays (40 μl) contained 10 μl membranes, ∽16 μl cytosol + homogenization bufffer (2 mg/ml final concentration), 4 μl 10X ATP regenerating system, 8 μl incubation buffer, 1 μl 10 μM synthetic miRNA (Integrated DNA Technologies, Coralville, IA) and 1 μl RNAsin (Promega). Reactions were incubated for 20 min at 30°C then placed on ice and mixed with 4.3 μl of 10X NEB buffer 3 and 1 μl of RNase I_f_ (50,000 units/ml) NEB, Ipswich, MA) was added to all reactions except to a no RNase control. Reactions were then incubated at 30°C for a further 20 min. Following incubation, RNA was immediately extracted according to Direct-Zol (Zymo Research) manufacturer’s instructions. First strand complementary DNA synthesis and quantitative PCR was performed using TaqMan miRNA assays (Life Technologies) for hsa-miR-223 or hsa-miR-190. Percent protection was calculated from the qPCR data by comparing the Ct of miRNA in the RNase treated samples against the no RNase control reaction (2^^^-(Ct_experimental_-Ct_control_)) in which the no RNase control was set to 100 percent.

#### Streptavidin pull-down of miR-223 and interacting proteins

The *in vitro* packaging assay was performed as described above with miR-223 with biotin linked to the 3’ phosphate (Integrated DNA Technologies). Samples were heated to 65°C for 20 min to inactivate RNase I_f_ and then mixed with 4.4 ul 10% Triton X-100 for a final concentration of 1% and kept on ice for 30 min. Novagen MagPrep Streptavidin-coated beads (10 μl/reaction) (EMD Millipore) were washed 3 times with 1 ml PBS and then added to the reaction lysate. The suspension was mixed by rotation for 2 h at 4°C, the beads were immobilized using a magnet and washed 3 times with 1 ml PBS. Proteins were eluted from bead-bound miR-223 with 50 μl 1 M KCl. In-solution liquid chromatography and mass spectrometry were performed according to standard procedures by the Vincent J. Coates Proteomics/Mass Spectroscopy laboratory (UC Berkeley).

#### Small RNA sequencing of cellular and exosomal RNA

RNA was prepared from cells and 3 l of HEK293T conditioned media. Sequencing libraries were generated using the Scriptminer Small RNA sequencing kit (Epicentre Biotechnologies, Madison, WI) from 1 μg total RNA from cells and 200 ng total RNA from exosomes according to the manufacturer’s protocol. The libraries were amplified and index barcodes were added by 11 cycles of PCR. Libraries were sequenced by 50 bp single read massively parallel sequencing on an Illumina Hi-Seq 2000 System at the Vincent J. Coates Genomic Sequencing Laboratory (UC Berkeley).

#### Small RNA sequencing analysis

Preprocessing of the 50 base pair single reads was filtered for read quality (read quality >20 and percent bases in sequence that must have quality >90) and adaptor sequences were clipped using the FASTX toolkit (http://hannonlab.cshl.edu/fastx_toolkit/) implementation on the GALAXY platform (usegalaxy.org) ((Blankenberg et al., 2010; Giardine et al., 2005; Goecks, Nekrutenko, Taylor, & Galaxy, 2010). Sequences were then mapped to the human genome (UCSC hg19 canonical build, UC Santa Cruz) with Bowtie2 using default settings (Langmead & Salzberg, 2012). Reads mapping to miRNAs were extracted from the dataset and normalized by dividing the number of reads mapping to each miRNA by the number of total reads mapping to all miRNAs and the quotient was then multiplied by one million (reads per million miRNA mapped reads – RPM). To analyze miRNA species, we used the quantifier program of the miRdeep2 software suite (Friedlander, Mackowiak, Li, Chen, & Rajewsky, 2012). Precursor only reads were determined by subtracting the number of reads mapping to mature (either targeting or passenger strand sequences) from the total number of reads mapping to the full-length precursor transcript for each miRNA. Those miRNAs with described passenger strands (star strand) were then analyzed to determine how many mature reads mapped to either targeting or passenger strands.

#### CRISPR/Cas9 genome editing

A pX330-based plasmid expressing venus fluorescent protein was kindly provided by Robert Tjian (Cong et al., 2013). A CRISPR guide RNA targeting the first exon of the YBX1 open reading frame was selected using the CRISPR design tool (Hsu et al., 2013). The YBX1 guide RNA was introduced into pX330-Venus by oligonucleotide cloning as described (Cong et al., 2013). HEK293T cells were transfected for 48 h at low passage number, trypsinized and sorted for single, venus positive cells in a 96 well plate using a BD Influx cell sorter. Wells containing single clones (16 clones) were allowed to expand and were screened by semi-nested PCR using primers targeting the genomic region flanking the guide RNA site. Primers for the first round of PCR (10 cycles) were: YBX1-F1 (GGTTGTAGGTCGACTGAATTA) and YBX1-R1 (ACCGATGACCTTCTTGTCC) The PCR primers from the first round were removed using DNA clean and concentrator-5 kit (Zymo Research) according to manufacturer’s instructions and the second round of PCR (25 cycles) was performed with primers: YBX1-F2 (CGGCCTAGTTACCATCACA) and YBX1-R1 (ACCGATGACCTTCTTGTCC). PCR products were separated on a 2.5% agarose gel to identify products smaller than the wild type PCR product, indicating a deletion. Clones (8) showing homozygous or heterozygous mutations were then screened by immunoblot to identify those that did not express YBX1. A clone containing a single homozygous mutation at the target site and not expressing YBX1 by immunoblot was recovered and designated ΔYBX1.

## Acknowledgements

We thank Amita Gorur for help and advice on electron microscopy. We also thank the staff at the UC-Berkeley shared facilities used in this work, the Functional Genomics Laboratory, the Vincent J. Coates Sequencing Facility, the Vincent J. Coates Proteomics Facility, the Computational Genomics Resource Laboratory and the Flow Cytometry Facility.

### Competing Interests

The authors declare no competing interests.

---

**Figure 1-figure supplement 1:**
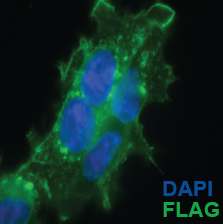
Sub-cellular localization of C-terminal CD63-luciferase-FLAG fusion.

